# The accuracy of NMR protein structures in the Protein Data Bank

**DOI:** 10.1101/2021.04.05.438442

**Authors:** Nicholas J Fowler, Adnan Sljoka, Mike P Williamson

## Abstract

We recently described a method, ANSURR, for measuring the accuracy of NMR protein structures. It is based on comparing residue-specific measures of rigidity from backbone chemical shifts via the random coil index, and from structures. Here, we report the use of ANSURR to analyse NMR ensembles within the Protein Data Bank (PDB). NMR structures cover a wide range of accuracy, which improved over time until about 2005, since when accuracy has not improved. Most structures have accurate secondary structure, but are too floppy, particularly in loops. There is a need for more experimental restraints in loops. The best current accuracy measures are Ramachandran distribution and number of NOE restraints per residue. The precision of structure ensembles correlates with accuracy, as does the number of hydrogen bond restraints per residue. If a structure contains additional components (such as additional polypeptide chains or ligands), then their inclusion improves accuracy. Analysis of over 7000 PDB NMR ensembles is available via our website ansurr.com.

## Introduction

We recently described a method, ANSURR (Accuracy of NMR Structures using Random Coil Index and Rigidity), which characterises the accuracy of NMR protein structures^1^. The method compares two measures of local rigidity: the *random coil index* (RCI), which uses backbone chemical shifts to estimate the fraction of random coil by residue^2^, and *rigidity theory*, which takes the structure, converts it to a constraint graph, and calculates rigidity using a pebble-game algorithm, based on the program Floppy Inclusions and Rigid Substructure Topography (FIRST)^3^. These measures are compared directly after re-scaling of RCI to ensure comparability, to generate scores measuring the accuracy of the structure, as described below. In our description of the method^1^, we compared ANSURR scores to a range of other measures that might be expected to provide a measure of accuracy, and also used the scores to provide a preliminary assessment of how NMR structures compare to X-ray crystal structures. These comparisons were carried out on a manually curated set of structures. In this work, we have expanded the comparison to study the accuracy and quality of all NMR structures within the PDB that match our selection criteria.

Ways of measuring the accuracy of NMR structures have been investigated previously^4-12^, including by a validation task force set up by the PDB^13^. These measures can be divided into two groups: *geometrical tests* and *comparisons to input data*. The geometrical tests are the same as those used as validation measures by X-ray crystallography and electron microscopy, and include measures of the proportion of residues within allowed or disallowed regions of the Ramachandran plot, atomic clashes, and packing density. These are robust and reliable measures, but are aimed at testing whether the protein structure has geometry that matches that obtained from high-quality experimental structures, rather than whether it matches well to the NMR input data. The refinement of NMR structures relies heavily on the quality of geometrical terms and force fields; much more than does X-ray structure refinement, because NMR structure calculations have far fewer experimental restraints. It is therefore reasonable to expect that geometrical tests should provide a guide to the accuracy of NMR structures, and indeed this was shown in our preliminary comparisons, where we found that the single best existing predictor of accuracy was the Ramachandran distribution: either the proportion of residues in the allowed region or the proportion in the disallowed region, which are strongly related.

However, the more interesting set of tests are the comparisons to input data, since these should more clearly discriminate the accuracy of structures. Crystallographers use the *R* factor, which directly compares the experimentally determined electron density with the density predicted by the structure. NMR spectroscopists have no equivalent measure. The most obvious equivalent is violations of NOE restraints, or possibly the number of NOE restraints per residue. The biggest challenge in using these as measures of accuracy is that NOE restraints are several removes from any experimental measurement. The experimental measurement is the NOESY spectrum. To extract restraints from the NOESY spectrum, the peaks must be picked: a person or a computer algorithm must decide which peaks are noise or experimental artifacts, and which intensities are distorted by peak overlap or baseline problems. There has to be a conversion from peak intensity (height or volume – not the same thing, though both are problematic in different ways) to distance. Classically this is done using a strong/medium/weak classification^14^, which avoids problems arising from unknown amounts of internal motion in the protein, but is a rather subjective system. The distances arising from this process are then fed into the structure calculation. This is an iterative process, mainly because most chemical shifts can be assigned to more than one nucleus, and so most NOE peaks are ambiguous, in the sense that a peak in the NOESY spectrum often cannot be assigned to a unique pair of protons, but rather a set of possible pairs. Structure calculations therefore iteratively reduce the ambiguity of existing restraints and possibly modify or add to the list of restraints. There is also typically a process of checking and possibly removing restraints that are repeatedly violated in structure calculations^15^. There is no theoretical justification for such an action, other than the observation that some errors in NOE assignment are inevitable because of incomplete knowledge. A final problem is that the best determined NOEs are the least useful, because they are typically between protons that are within the same amino acid residue. Some practitioners often leave out such NOEs because they have no information content, while others leave them in. Most NOEs in multidimensional spectra occur in more than one location within the spectrum: practitioners differ in whether only one or both such occurrences are used. Thus even such a fundamental measure as the number of NOE restraints is not well defined. All of this means that comparisons to input data, such as NOE restraint violations or numbers of restraints per residue, are ill-defined and handled differently by different practitioners.

ANSURR provides a different type of validation, based on chemical shifts rather than distance restraints. Backbone chemical shifts can often be obtained almost automatically from spectra and are generally reliable. They are also easily available for any proteins with NMR structures, not least because it is a requirement of PDB that chemical shift assignments be deposited with BioMagResBank (BMRB) at the same time as structure deposition with PDB. This makes them good parameters for validation.

In this paper, we explore what ANSURR can tell us about the accuracy of NMR structures in the PDB, and conclude that although some structures are excellent, they vary considerably in their accuracy. Following a general survey of PDB structures, we look at the different measures of structure quality, to evaluate what they can tell us about structure quality. Finally, we look at the accuracy of oligomers and complexes, and show that inclusion of all molecular components is beneficial to the accuracy.

## Results

We noted previously^1^ that RCI is only reliable when the chemical shift completeness of backbone shifts (HN, N, Cα, Cβ, Hα, C’) is at least 75%. We have therefore used all PDB NMR structures for which a BMRB chemical shift assignment file exists at 75% completeness or more, and where the polypeptide chain is at least 20 residues long. This results in a subset of PDB structures, here named PDB75, which contains 4742 ensembles. Further details can be found in Methods.

ANSURR performs two different comparisons (Fig. 1): (1) The *correlation* between the rigidity computed from chemical shifts (RCI) and from the structure using rigidity theory (FIRST). This tests whether peaks and troughs in the two measures are in the same places. Because a peak is a locally flexible region (typically a loop) and a trough is a locally rigid region (typically a region of regular secondary structure), this measure is largely detecting whether secondary structure is correct. It is however rather more subtle than a simple comparison of locations of secondary structure in that it includes a comparison of breaks and weaknesses in secondary structure. (2) The root-mean-square-deviation (RMSD) between the RCI and FIRST outputs. This tests whether the overall rigidity of the structure matches the rigidity as defined by RCI. An important determinant of local rigidity is the presence of nonbonded interactions i.e. hydrogen bonds and hydrophobic contacts. This comparison is thus to a large extent a measure of whether the structure is close enough to the correct structure for nonbonded interactions to be formed correctly. The two comparisons measure different aspects of the structure and are therefore not strongly correlated.

**Figure 1.**
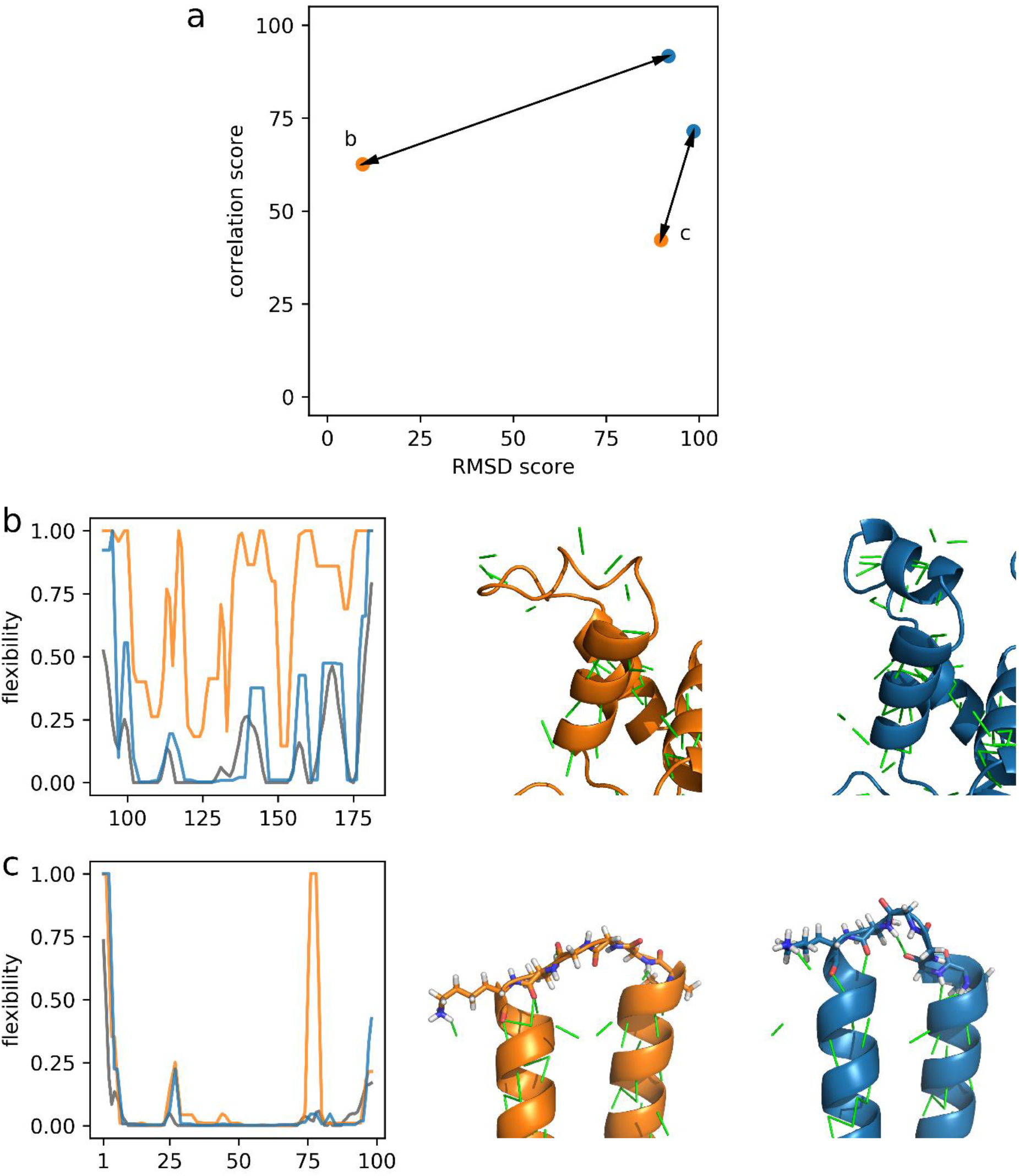
Outline of ANSURR output. (a) ANSURR compares measures of local rigidity produced from chemical shifts (RCI) and from the structure according to rigidity theory (FIRST). It comprises *correlation*, which measures whether peaks and troughs are in the same places; and *RMSD*, which compares overall rigidities. The raw values are converted into *scores*, which are rank percentile values relative to all NMR protein structures in the PDB75 dataset (see Figure 2). (b) Left: ANSURR analysis of model 17 of the DNA-binding domain of the human Forkhead transcription factor AFX (PDB ID 1e17) from the RECOORD^16^ CNS [refined in vacuo (orange)] and CNW [refined in explicit solvent (blue)] datasets. RCI values are in grey. The CNW structure is more accurate, as measured by both correlation and RMSD scores. Middle and right: Residues 120-140 of the CNS and CNW structures, respectively. Hydrogen bonds are indicated by green lines. Drastically improved ANSURR scores, mostly RMSD score, are observed following refinement in explicit solvent. This largely originates from the increased accuracy of nonbonded interactions (hydrogen bonds and hydrophobic contacts). (c) Left: ANSURR analysis of chain A from model 7 (orange) and model 2 (blue) of the designed protein XAA (PDB ID 6o0i). RCI values are in grey. The small loop between residues 74-79 in model 7 is too floppy but correctly rigid in model 2, resulting in a better correlation score for model 2 (a). Middle and right: The loop of models 7 and 2, respectively. The rigidity of the loop in model 2 is determined by a single hydrogen bond which is not present in model 7, suggesting that the loop in model 7 is not representative of the solution structure.

We have carried out ANSURR analyses of all 4742 NMR ensembles in the PDB75 dataset. These analyses are available from our website ansurr.com. In Fig. 2a we present the average correlation and RMSD values for each ensemble, shown as a two-dimensional plot. Poor structures are found in the lower left corner, while good structures are in the upper right corner of such a plot. Structures span a wide range of accuracy, with some very poor structures. Conversely, a significant number of structures are of very good accuracy. The most densely populated region has a Spearman’s rank correlation coefficient ρ of around 0.7, indicating that most NMR structures in PDB75 have essentially the correct secondary structure. The most common RMSD value is however only around 0.3, indicating that the structures have overall rigidity that does not match RCI well. In almost all cases, the rigidity of the structure is lower than the rigidity implied by RCI, ie NMR structures are too floppy. For most NMR structures, the rigidity in secondary structures is good, and the floppiness is mainly in the loops (compare the two-dimensional plots on the left of Fig. 1b,c, where the troughs have rigidities close to zero on both measures, and therefore match well, while the peaks are more variable). Those who determine NMR structures have tended to concentrate on secondary structure and not worried too much about loops, the general feeling being that loops are probably fairly undefined in solution anyway, so that the lack of definition in loops (ie the spread of structures within an ensemble) is probably “real”. Our analysis indicates that this is not true – structures in solution are considerably more structured and defined in loops than they are in typical NMR PDB structures. For example, compare the loops in Fig. 1c, where addition of a single hydrogen bond results in a large improvement in accuracy. We suggest that this represents a failing in many NMR structures.

**Figure 2.**
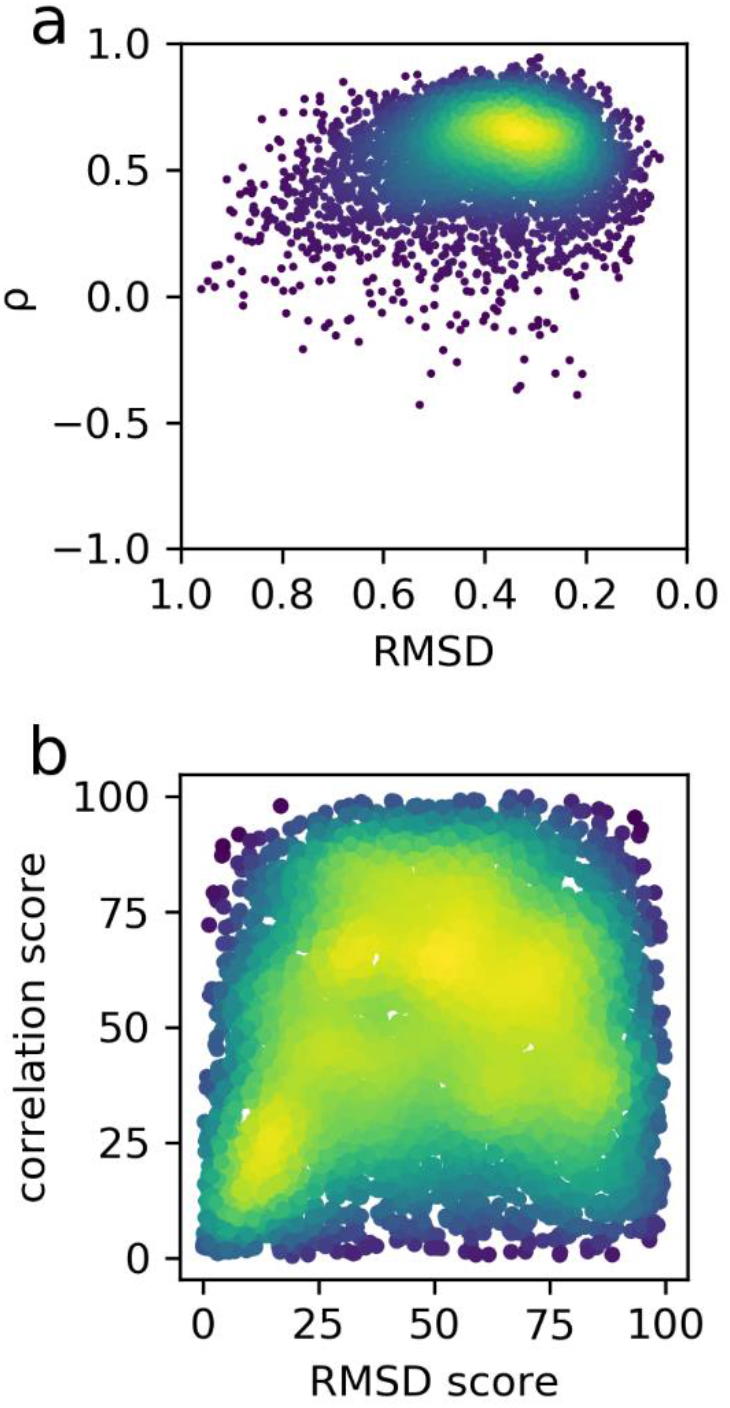
Distribution of ANSURR measures from the PDB75 dataset. For each ensemble in the dataset, we display the mean values for the ensemble rather than individual scores. (a) Raw values for correlation and RMSD. Note that that the RMSD values are displayed on a descending scale, for ease of comparison to scores. (b) The raw values have been convertible to percentile rank scores, which range from 0 (the worst structure in the PDB75) to 100 (the best). Colours represent the density of values.

The data in Fig 2a are nonlinear, making it difficult to judge by how much any individual ensemble is better or worse than typical PDB results. An alternative presentation of these data is thus to show the results as a percentile of the complete PDB75 dataset. We term these percentile values the correlation score and RMSD score, which by definition run from 0 to 100, and are summarised in Fig. 2b. An advantage of the percentile scores is that the distributions of RMSD and correlation scores are more comparable, meaning that the correlation and RMSD scores can be summed to give an overall ANSURR quality score that measures the overall structural accuracy reasonably well, whereas the same cannot be said of the correlation and RMSD values themselves. In the subsequent analysis, we therefore add the correlation and RMSD scores together to produce a single accuracy score, termed ANSURR score, when it makes sense to use a single overall measure. The individual results for RMSD and correlation scores are shown in supplementary information.

### Trends in quality

In our previous study^1^ we noted that a critical factor in improving accuracy was refinement in explicit solvent, which increased the ANSURR score by approximately 35. In this study we were unable to carry out such a comparison, because we were unable to work out, either from the PDB header or even from original publications in many cases, whether and how such refinement had been carried out. This further highlights the importance of reporting guidelines for PDB depositions. As a proxy, we looked at the relationship between ANSURR score and year of deposition (Fig 3a).

**Figure 3.**
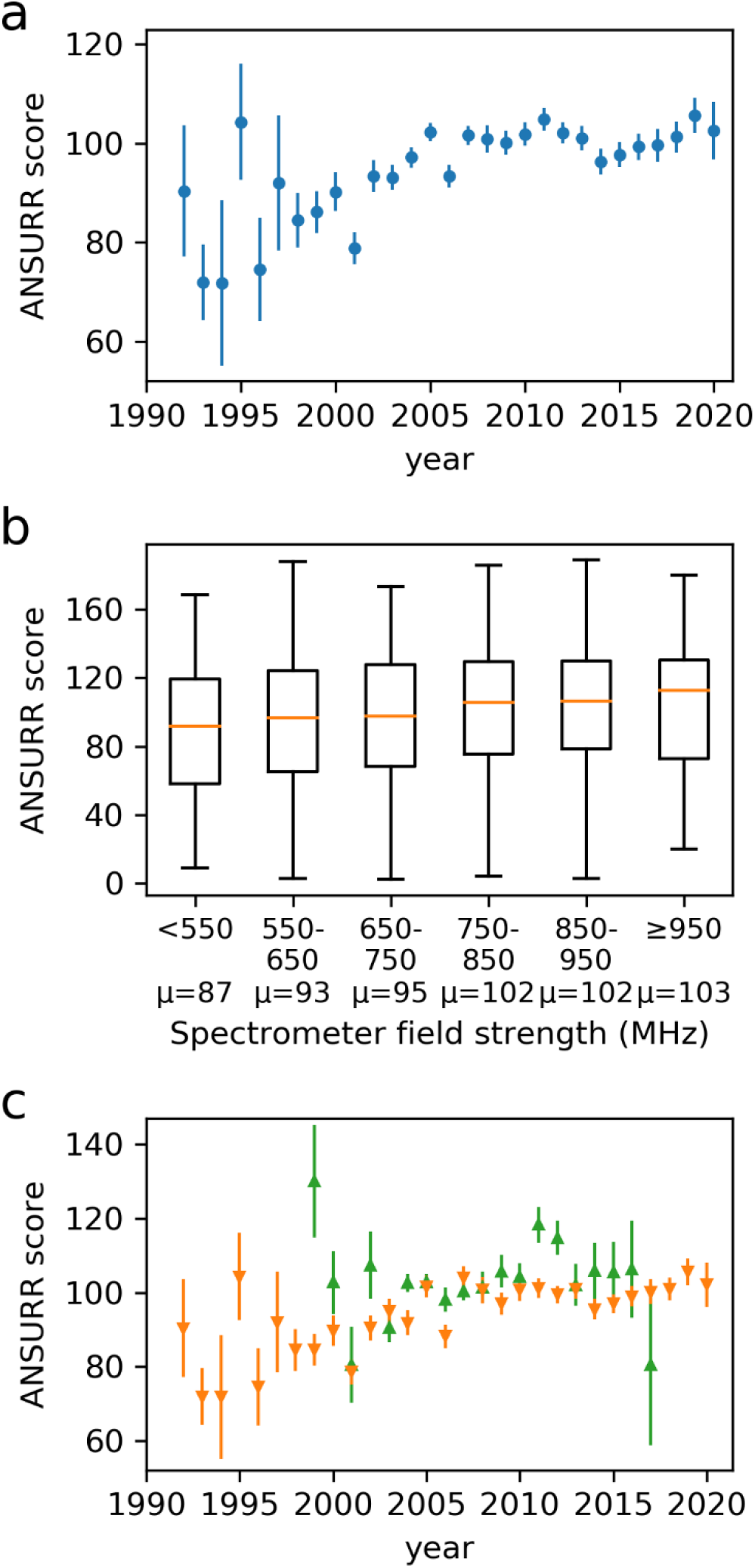
Trends in NMR accuracy with time. (a) ANSURR score (sum of correlation and RMSD scores) for PDB75 NMR ensembles, as a function of year of deposition. Data are mean ± standard error of the mean. Data points are plotted only for years with at least 3 ensembles from the PDB75 dataset. (b) ANSURR score *vs* highest field strength cited in the PDB header. Data are shown as box plots, and indicate the median (orange lines), first and third quartiles (box) and extremes. Mean values are indicated below the plot. (c) ANSURR score *vs* year of deposition. This shows the same data as in (a) but split into structures coming from structural genomics consortia (green) and all others (orange). Sample sizes for each plot are provided in supplementary information (SI Table 1).

It is clear that there was a gradual increase in accuracy with time, up to about 2005, after which there is little obvious improvement. It is probably significant that the key papers on refinement in explicit solvent were published in 2003^17,18^, and many of the standard methods for calculating NMR structures (eg TALOS for dihedral restraints^19^, CYANA^20^, XPLOR-NIH^21^) were also established in 2003 or shortly before. Since then, there have been no step changes in the way protein NMR structures were calculated. We hope that the results shown here will stimulate interest in improving the quality of NMR structures.

There are several confounding factors that may also contribute to the improvement in structural accuracy with time. One is the steady improvement in spectrometer sensitivity, largely from increasing commercially available field strength, which will improve structural accuracy by providing higher sensitivity and thus more complete NOE restraints. The results of this analysis are shown in Fig 3b, which show that field strength has a significant effect on accuracy.

We also looked to see if specific research groups did noticeably well or badly in terms of accuracy. There are too few structures from any individual group to produce useful conclusions, but we were able to compare the accuracy of structures originating from structural genomics consortia to all other structures (consortia are listed in SI Table 2). The results are shown in Fig 3c, and suggest that structural genomics consortia have tended to generate slightly more accurate structures, though the differences are small. It is worth noting that structural genomics consortia have also tended to focus on “low-hanging fruit”, which yield good NMR spectra, and therefore might be expected to be more accurate for that reason.

The trends with time are more apparent if we consider ANSURR scores obtained for all structures regardless of backbone chemical shift completeness (SI Fig2). One would hope that this is because we have better statistics with more structures, but these results must be interpreted carefully as the accuracy of RCI is less reliable when chemical shift completeness is below 75%.

### Comparison to established data-driven quality measures

In our earlier study^1^ we looked at the correlation between ANSURR scores and a range of quality measures typically used to characterise NMR protein structures. In that study we used a set of 173 ensembles that had been processed and refined in a consistent way, with manually curated checks on input restraints^16^. Here we carry out a similar analysis, but using all ensembles from the PDB75 dataset. It is a much larger dataset, but also a much less uniform one. For simplicity, we present here summed ANSURR scores, discussing the component correlation and RMSD scores where relevant. RMSD and correlation scores plotted separately are provided in supplementary information (SI Fig3).

There is a strong relationship between ANSURR score and the number of NOE restraints per residue (Fig 4a). We noted in the Introduction that there is a lack of consistency among NMR practitioners in the counting of NOE restraints, but this does not hide the expected influence of number of NOEs on accuracy. The effect is seen almost entirely on RMSD score rather than correlation score (see SI Fig3a); in other words, increased numbers of NOE restraints help tie down local structure better, rather than improving the definition of secondary structure.

**Figure 4.**
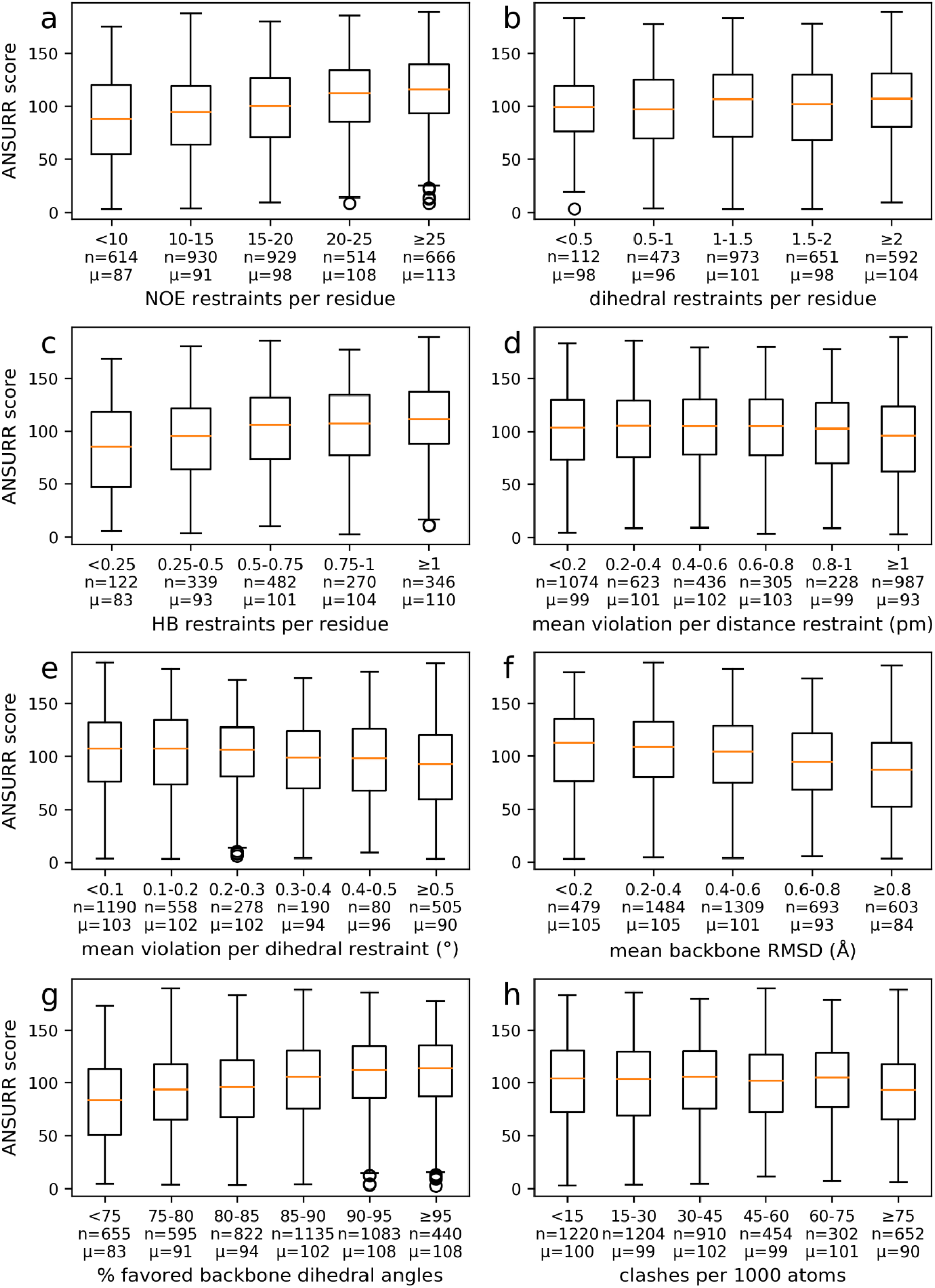
Dependence of ANSURR scores on other measures of structural accuracy. Data are presented as in Fig 3b, as box plots. The number of samples and the mean are indicated below each box. (a) Number of NOE restraints per residue. (b) Number of dihedral restraints per residue. (c) Number of hydrogen bond restraints per residue. (d) Mean size of distance restraint violation (Å). (e) Mean size of dihedral restraint violation (°). (f) Mean backbone root-mean-square-difference (RMSD – the precision). (g) Percentage of backbone (φ, Ψ) pairs within the favoured regions of the Ramachandran plot, as measured by MolProbity. (h) Clashscore (clashes per 1000 atoms), as measured by MolProbity.

By contrast, the relationship between the number of dihedral restraints per residue and ANSURR score is less apparent (Fig 4b). Nonetheless, structures with at least two dihedral restraints per residue (mean ANSURR score of 104) tend to score better than those with zero (mean ANSURR score of 97, two-sided *t*-test *p*-value = 1.4 ×10^−5^). If we consider RMSD and correlation scores separately (SI Fig3b), we see that RMSD score increases with the density of dihedral restraints, but at the expense of correlation score. This suggests that dihedral restraints act to rigidify a structure by improving backbone geometry but are perhaps too weak to improve the overall accuracy significantly.

There is a strong relationship between ANSURR score and the number of hydrogen bond restraints per residue (Fig 4c). This relationship arises entirely from the RMSD score (SI Fig3c), and shows that NMR structures have greatly improved rigidity when hydrogen bond restraints are present. This effect is most noticeable in loops, which in NMR structures are generally too floppy in comparison to the RCI values. In crystal structures, there are usually hydrogen bonds that serve to rigidify the structure, and our results indicate that such hydrogen bonds are retained in solution, since adding them produces structures that match well to RCI values (to be published). In the absence of hydrogen bond restraints, much of the hydrogen bonding network will be determined by the forcefield used during refinement. However, the forcefield alone is clearly unable to induce hydrogen bonds when they are not present in the unrefined structure. There is thus a clear implication: NMR structures will be greatly improved in their overall accuracy (particularly in the loops) if we can find experimental restraints that define the hydrogen bonds. To date the only reliable hydrogen bond restraint comes from ^3h^*J*_NC’_ scalar couplings, usually obtained from long-range HNCO spectra^22^. Such couplings are small, and typically only observable for small proteins with long relaxation times. There is therefore a challenge for NMR, to identify suitable hydrogen bond restraints that can be used to improve structures.

Neither distance nor angle violations particularly correlate with ANSURR score, although structures with higher level of violations (>0.01 Å per distance restraint, >0.3° per dihedral restraint) tend to score slightly worse (Fig 4d, e). This suggests that violations can help to diagnose structures with significant errors but may not be a useful guide to accuracy. It is worth repeating the comments made above, that in many structure calculations, restraints that are repeatedly violated are omitted or modified in subsequent iterations, implying that violation statistics may not be reliable guides to accuracy^23-25^.

There is a moderate relationship between the ANSURR score and the backbone RMSD between structures in the ensemble (the precision of the ensemble; Fig 4f). In other words, the accuracy and precision of the ensemble are related: more accurate ensembles also have tighter precision. This finding is different from what we saw previously with a manually curated set of structures^1^, where there was very little correlation between accuracy and precision. We hypothesise that in structure calculations where the NOE network is sufficiently dense to limit the precision effectively, the NOEs will also tend to improve the accuracy, but this merits further investigation.

### Comparison to geometrical tests

There is a clear relationship between ANSURR score and the percentage of residues in the favoured region of the Ramachandran plot (Fig 4g). The process of NMR structure determination is a joint refinement against both the experimental restraints and knowledge-based parameters (van der Waals packing, coulombic forces, bond/angle potentials, solvent interactions etc) and it is therefore not a surprise that accurate structures should also have good Ramachandran distributions. On the other hand, there is very little relationship with clashscore (Fig 4h). We saw a similar effect in our previous study^1^.

### Biological assemblies and ligands

About 13% of NMR structures in the PDB comprise more than one polymer chain, ie are oligomers or have bound peptides. It has been previously shown using X-ray crystal structures that interactions between subchains can significantly affect the rigidity of biological assemblies as a whole^26^. Here we analyse ANSURR scores computed for 550 biological assemblies that were calculated using NMR). Figure 5a shows the difference between the ANSURR score obtained when rigidity is computed for the entire biological assembly and when rigidity is computed for subchains individually. ANSURR score improves for 92% (504/550) of NMR ensembles when rigidity is computed for the entire biological assembly. The results clearly demonstrate that the formation of biological assemblies significantly affects rigidity and in a way that leads to better agreement with rigidity calculated from chemical shifts. It might be expected that greater improvements in ANSURR score would be seen for assemblies which share a larger degree of contact between the constituent chains, and this is indeed the case. In figure 5b the ratio of the solvent accessible surface area (SASA) summed over individual chains and that for the biological assembly (assemblies with larger degree of contact between chains will have greater SASA ratios) is plotted against the difference in ANSURR score. As an example, figure 5c shows the oligomer 2MJA for which the ANSURR score increases by 90 when rigidity is computed for the entire assembly. It is easy to see why. The structure is made up of beta strands formed by a combination of the two chains, which when separated would become considerably more flexible^27^.

**Figure 5.**
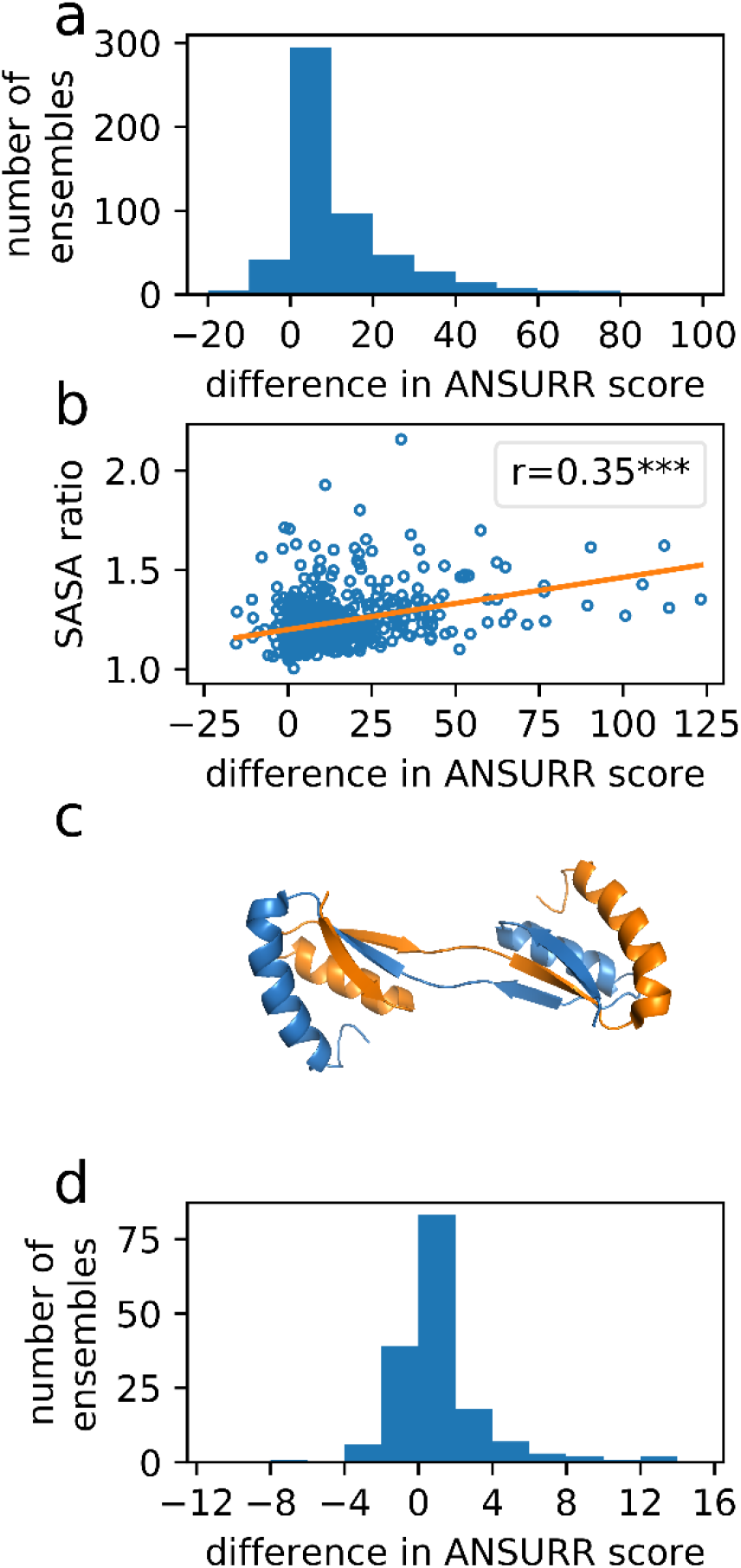
The effect of including other components in a biological assembly in the PDB75 dataset. (a) The change in ANSURR score when the entire multichain assembly is used for the ANSURR calculation, as opposed to calculating the scores for subchains independently. The mean change is 12. (b) Weak but significant correlation (Pearson’s r=0.35, two-tailed p-value=7.8×10^−18^) between the ratio of solvent accessible surface area summed over individual chains and that for the biological assembly, and the difference in ANSURR score. (c) The structure of the protease GlpG (PDB 2MJA), which comprises a domain-swapped dimer, and has a large improvement in ANSURR score when rigidity is calculated for the dimer, compared to the two monomers separately. (d) The change in ANSURR score when adding non-peptide partners to the assembly, such as small molecule ligands and metals. The mean change is 1.

About 15% of NMR protein structures contain non-polymer instances e.g. ligands such as drug molecules and metals. We analysed ANSURR scores computed for 162 NMR ensembles with bound ligands. Figure 5d shows the difference in ANSURR score obtained when rigidity is computed with ligands present and without. ANSURR score improves for 57% of the ensembles, is unchanged for 15% of ensembles and is worse for 28% of ensembles. Overall, the total change in ANSURR score is much less than in our biological assemblies analysis above. This is to be expected as bound ligands are generally much smaller than polymer chains in biological assembles and would be expected to make much less difference to the total flexibility. Regardless, it seems that more often than not, bound ligands do alter the rigidity of the structure, and in a way that tends to improve agreement with rigidity calculated from chemical shifts.

## Discussion

We report on the application of a new method, ANSURR, which measures the accuracy of NMR protein structures. Previous methods for assessment of accuracy have relied either on analysis of NOE violations, or on the precision of the ensemble. Both of these are poorer measures of accuracy, for reasons discussed here. Our method compares the rigidity of the structure to the rigidity indicated using a modified version of the random coil index, and thus provides a measure of accuracy that compares experimental measurements to the final structure. The method has been applied to all NMR ensembles in the PDB that have chemical shift data in BMRB. The analysis presented here has focused on ensembles with at least 75% backbone chemical shift completeness and at least 20 amino acid residues, but can be applied to any NMR structure. Our website ansurr.com provides details for all PDB NMR structures with chemical shifts in the BMRB, with warnings for those ensembles calculated with less than 75% chemical shift completeness.

ANSURR scores can be calculated rapidly and easily for any NMR protein structure, and (with the help of the underlying data, Fig. 1) provide a simple and user-friendly measure of accuracy. The PDB currently provides a slider bar assessment of structure quality for each deposited ensemble, but this focuses on geometrical quality rather than accuracy. ANSURR provides a general measure of accuracy for NMR protein structures, which we hope will be useful for the structural biology community. A majority of the users of PDB are not depositors, and are thus not ‘structural biology experts’ but scientists looking for the insight that can be provided by structural details. For such users, NMR structures have been problematic because it has been unclear how accurate they are, and therefore whether NMR structures can be used with the same confidence that (for example) crystal structures can. ANSURR goes at least some way to answering this problem, by providing a measure of accuracy that does not rely on NOE restraints.

The analysis provided here shows that there is a large range in the accuracy of NMR structures in the PDB. Structure calculation improved steadily up to about 2005, since when there has been little change in overall accuracy. The majority of structures have good correlation between rigidity calculated from chemical shifts and from the structure, indicating that the regular secondary structure is generally correct. Our previous paper^1^ made a comparison between NMR structures and crystal structures, within a limited number of curated structures, and showed that from correlation scores, the secondary structure of NMR and crystal structures are of comparable accuracy.

However, the same cannot be said for RMSD score. Our analysis shows that the large majority of NMR structures are too floppy, by comparison to the “true” rigidity indicated by RCI. This applies throughout the structures, but because regular secondary structure is inherently fairly rigid, it is more evident, and more troubling, in loops. Structural biologists tend to compare structures by overlaying them for best fit over backbone atoms, and then displaying them as cartoon plots, which emphasise the locations and orientation of regular secondary structure elements. This is a sensible practice, but it leads to the widespread assumption that the important features of a protein are its regular secondary structure, and that loops are relatively unimportant. Indeed, at least among the NMR community, there is a general feeling that loops in solution are probably not well defined, and that the variability seen in loops (and differences in loops between NMR structures and crystal structures) reflect the “real” flexibility of loops and are not a major cause for concern. Our analysis shows this is not true: most loops in solution are much less flexible than is indicated by the ensemble, and loops in NMR structures are for the most part underdetermined and inaccurate. A recent publication^28^ reaches similar conclusions, using different methodology. This is a consequence of the fact that there are usually very few NOE restraints in loops, and often very few restraints at all. Our analysis implies that NMR spectroscopists need to work harder to identify structural restraints within loops, because this will significantly improve the overall accuracy of the structures. Methods for the identification of hydrogen bonds would be of particular importance, because of the power of hydrogen bonds to limit flexibility.

Here, we have compared ANSURR scores to a range of parameters that might be considered to provide a measure of accuracy. We have shown that the distribution of backbone dihedral angles within the Ramachandran surface is a good measure of accuracy (Fig. 4g), as are the number of NOE restraints (Fig. 4a) and the number of hydrogen bond restraints (Fig. 4c). We tried to make comparisons with other factors, in particular the method of structure refinement (for example, inclusion of explicit solvent) and the programs and parameters used for structure calculation and refinement, but were unable to do so, because the PDB record fields do not hold this information in a consistent way, and we were unable to come up with a machine-readable way of getting the information. Indeed in many cases it is not possible to be confident what the authors did even from reading the relevant papers. There is thus a need to generate more consistent and machine-readable documentation of the methodology used to calculate NMR structures. We note that PDB and BMRB are discussing the introduction of a NMR Exchange Format which could help to improve such documentation^29^.

The results presented here make it clear that there remains considerable scope for improving the accuracy of NMR structures, particularly in loops, which is where most ligand binding sites and enzyme active sites are located^30,31^. Better measures of accuracy are likely to drive better accuracy, which can only benefit the entire community.

## Methods

### ANSURR calculations

7187 NMR protein structure ensembles and their corresponding backbone chemical shifts were downloaded from the PDB and BMRB, respectively. Paired PDB and BMRB IDs are provided as a supplementary text file (pdb_chain_bmrb.txt). ANSURR was used to validate each model in each NMR ensemble with the following options: re-reference chemical shifts using PANAV, include non-standard residues, include ligands and combine chains when computing flexibility. As the number of models in an ensemble varies (from a single model to hundreds), we averaged the correlation, RMSD and ANSURR scores for all members of each ensemble. Scores for each model are provided as a supplementary text file (ansurr_scores_Nov2020.out) and can also be downloaded from our website ansurr.com. Unless stated otherwise, the analysis in this work was performed on the subset of ensembles with at least 20 residues and at least 75% chemical shift completeness. This subset is termed PDB75 and comprises 4742 ensembles.

### Established data-driven quality measures

The quality measures used in our analysis presented in figure 4 were generated as follows. The number of restraints per residue and the mean restraint violations per residue were acquired from the NMR Restraints Grid (restraintsgrid.bmrb.wisc.edu)^32,33^. Mean backbone RMSDs were extracted from PDB validation reports for each ensemble with more than 1 model. The average percentage of favoured backbone dihedral angles was computed for each ensemble using the program ramalyze, part of the Molprobity suite^34^. The program clashscore (also part of Molprobity) was used to compute the average number of clashes per 1000 atoms for each ensemble.

### Dataset of oligomeric NMR protein structures

The PDB was searched for oligomeric NMR protein structures using the advanced search function on the RCSB PDB website (www.rcsb.org). This set of structures was then filtered to include only those which had a corresponding set of backbone chemical shifts on the BMRB with at least 75% chemical shift completeness and had at least 20 residues in each chain. The final set comprised 550 protein NMR structures. ANSURR scores when rigidity is computed for the entire biological assembly are provided in ansurr_scores_Nov2020.out. ANSURR scores when rigidity is computed separately for each subchain are provided as a supplementary text file (ansurr_scores_asmonomers.txt)

### Dataset of NMR protein structures containing free ligands

The advanced search function on the RCSB PDB website was used to obtain NMR protein structures that contained at least one non-polymer instance. An in-house program was used to exclude structures with ligands which contained metals because the strength of bonds which include metals cannot be determined from a protein structure without further calculations e.g. with density functional theory or quantum mechanics. This set was then filtered to include only those which had a corresponding set of backbone chemical shifts in the BMRB with at least 75% chemical shift completeness and had at least 20 residues in each chain. The final set comprised 162 protein NMR structures. ANSURR scores when rigidity is computed with ligands included are provided in ansurr_scores_Nov2020.out. ANSURR scores when rigidity is computed without ligands included are provided as a supplementary text file (ansurr_scores_noligands.txt)

## Acknowledgements

We thank the Biotechnology and Biological Science Research Council (BBSRC) for funding to N. J. F. (BB/P020038/1), and CREST, Japan Science and Technology Agency (JST) and PRISM for funding to A. S.

## Data availability

All study data are provided in supplementary information. In addition, ANSURR scores for 7187 NMR protein structures can be viewed and downloaded from our website ansurr.com.

## Author contributions

N.J.F. performed research; all authors designed research, analysed data, and wrote the paper.

## Data deposition

The ANSURR program and associated documentation can be downloaded from github.com/nickjf/ANSURR, https://doi.org/10.5281/zenodo.416158660. A typical calculation on an ensemble of 20 models for a 150-residue protein takes less than a minute. ANSURR output for all PDB ensembles can be found on ansurr.com.

## Additional information

Supplementary information is available.

