## Supplementary Information for "The accuracy of NMR protein structures in the Protein Data Bank"

Nicolas J Fowler[1], Adnan Sljoka[2,3,] and Mike P Williamson[1,*]

[1]Dept of Molecular Biology and Biotechnology, University of Sheffield, UK; [2]RIKEN Center for Advanced Intelligence Project, RIKEN, 1-4-1 Nihombashi, Chuo-ku, Tokyo, 103-0027 Japan; [3]Dept of Chemistry, University of Toronto, UTM, 3359 Mississauga Road North, Mississauga, ON, L5L 1C6, Canada

**Supplementary Figure 1.** Average RMSD, correlation and ANSURR scores for all structures in the PDB75 dataset according to secondary structure content. Scores for helical proteins (α) tend to be better than those for beta-sheet-rich proteins (β). Proteins with mixed content (αβ) have scores in between. We noted a similar trend in our previous study of a curated dataset (Fowler et al. A method for validating the accuracy of NMR protein structures. *Nature Communications* **11**, 6321 (2020)).

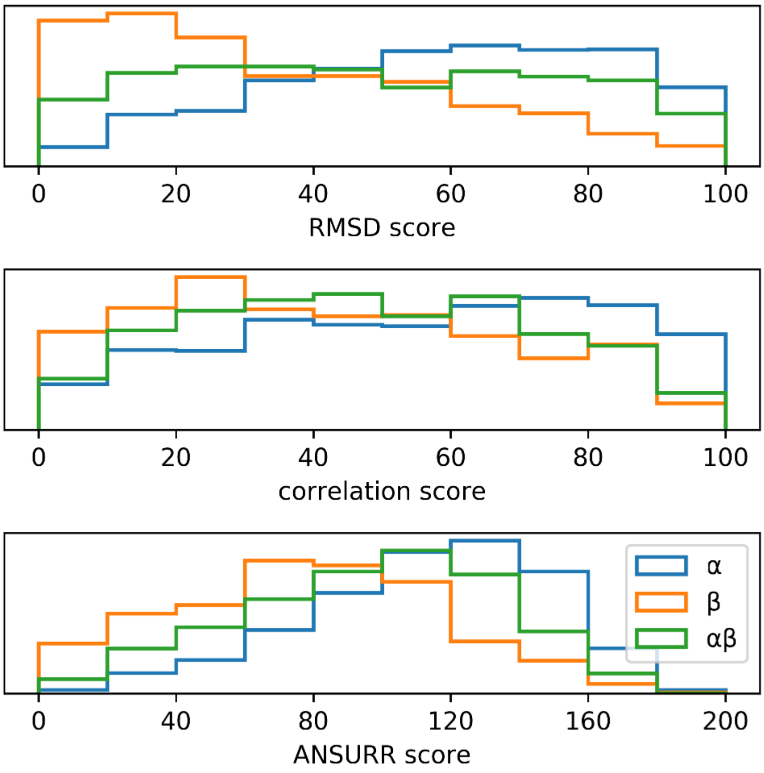

**Supplementary Table 1.** Sample sizes for Figure 3.

| | | |
|---|---|---|
| **Fig3a** | 1992 3, 1993 3, 1994 6, 1995 8, 1996 13, 1997 11, 1998 51, 1999 85, 2000 86, 2001 123, 2002 117, 2003 197, 2004 265, 2005 343, 2006 267, 2007 338, 2008 178, 2009 235, 2010 242, 2011 303, 2012 329, 2013 300, 2014 239, 2015 270, 2016 229, 2017 192, 2018 152, 2019 118, 2020 39 | |
| **Fig3b** | <550 164, 550-650 1145, 650-750 358, 750-850 2214, 850-950 696, ≥950 90 | |
| **Fig3c** | SGC<br><br>1999 3, 2000 4, 2001 11, 2002 20, 2003 79, 2004 132, 2005 196, 2006 138, 2007 214, 2008 66, 2009 83, 2010 82, 2011 65, 2012 57, 2013 46, 2014 20, 2015 24, 2016 10, 2017 6 | Non-SGC<br><br>1992 3, 1993 3, 1994 6, 1995 8, 1996 13, 1997 11, 1998 50, 1999 82, 2000 82, 2001 112, 2002 97, 2003 118, 2004 133, 2005 147, 2006 129, 2007 124, 2008 112, 2009 152, 2010 160, 2011 238, 2012 272, 2013 254, 2014 219, 2015 246, 2016 219, 2017 186, 2018 150, 2019 117, 2020 38 |

**Supplementary Table 2.** List of structural genomic consortia (SGC) with structures in the PDB75 dataset. Note that some structures were jointly produced by multiple SGC.

| **Structural genomics consortium** | **# of structures** |
|---|---|
| RIKEN Structural Genomics/Proteomics Initiative (RSGI) | 571 |
| Northeast Structural Genomics Consortium (NESG) | 476 |
| Joint Center for Structural Genomics (JCSG) | 82 |
| Center for Eukaryotic Structural Genomics (CESG) | 53 |
| Seattle Structural Genomics Center for Infectious Disease (SSGCID) | 40 |
| Structural Genomics Consortium (SGC) | 34 |
| Ontario Centre for Structural Proteomics (OCSP) | 34 |
| Structural Proteomics in Europe (SPINE) | 28 |
| New York Structural Genomics Research Consortium (NYSGRC) | 13 |
| Partnership for T-Cell Biology (TCELL) | 13 |
| Berkeley Structural Genomics Center (BSGC) | 10 |
| Structure 2 Function Project (S2F) | 9 |
| Center for Structural Genomics of Infectious Diseases (CSGID) | 7 |
| Midwest Center for Structural Genomics (MCSG) | 6 |
| Partnership for Stem Cell Biology (STEMCELL) | 6 |
| Mitochondrial Protein Partnership (MPP) | 6 |
| Montreal-Kingston Bacterial Structural Genomics Initiative (BSGI) | 5 |
| Chaperone-Enabled Studies of Epigenetic Regulation Enzymes (CEBS) | 5 |
| New York SGX Research Center for Structural Genomics (NYSGXRC) | 4 |
| TB Structural Genomics Consortium (TBSGC) | 1 |
| Membrane Protein Structures by Solution NMR (MPSbyNMR) | 1 |
| Center for Structures of Membrane Proteins (CSMP) | 1 |
| Assembly, Dynamics and Evolution of Cell-Cell and Cell-Matrix Adhesions (CELLMAT) | 1 |

**Supplementary Figure 2.** Trends in NMR accuracy with time. (a) ANSURR score (sum of correlation and RMSD scores) for 6703 NMR ensembles, as a function of year of deposition. Data are mean ± standard error of the mean. Data points are plotted only for years with at least 3 ensembles. (b) ANSURR score vs highest field strength cited in the PDB header. (c) ANSURR score vs year of deposition. This shows the same data as in (a) but split into structures coming from structural genomics consortia (green) and all others (orange). Sample sizes for each plot are provided in SI Table 3. Consortia are listed in SI table 4.

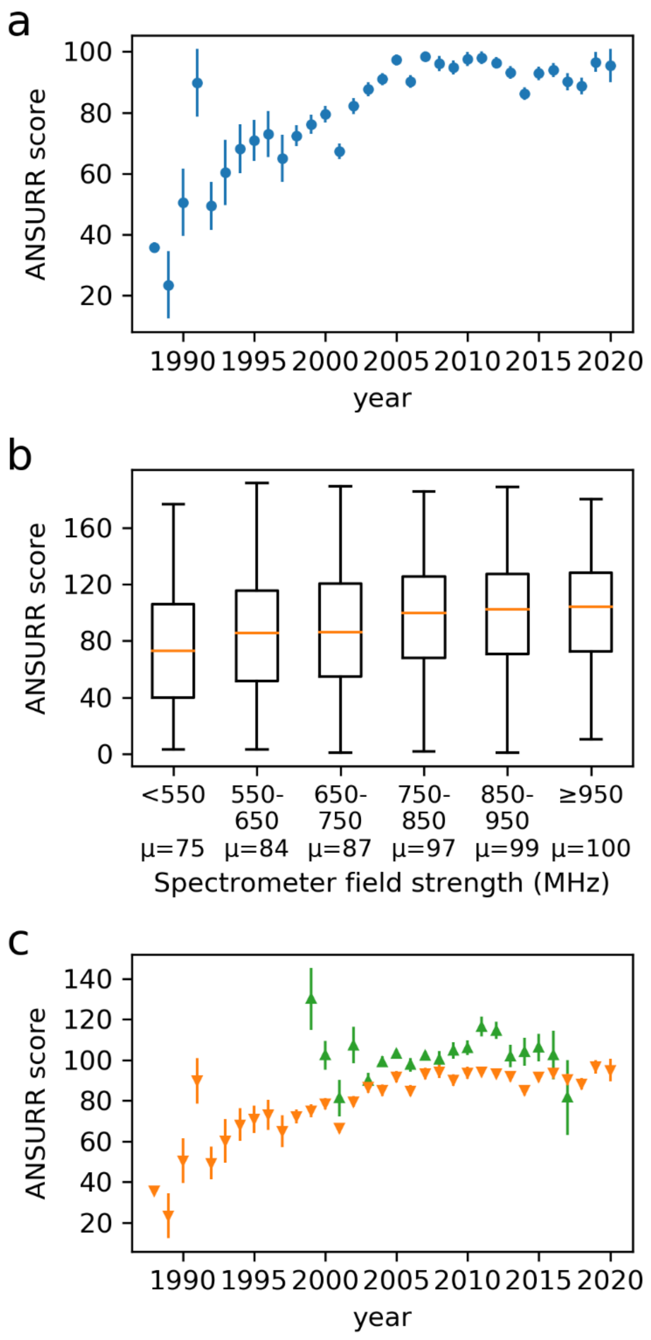

**Supplementary Table 3.** Sample sizes for SI Figure 2.

| | | |
|---|---|---|
| **SI Figure 2a** | 1988 3, 1989 3, 1990 12, 1991 11, 1992 20, 1993 16, 1994 15, 1995 34, 1996 23, 1997 29, 1998 117, 1999 148, 2000 164, 2001 220, 2002 214, 2003 270, 2004 342, 2005 409, 2006 349, 2007 444, 2008 233, 2009 290, 2010 297, 2011 407, 2012 434, 2013 409, 2014 363, 2015 360, 2016 327, 2017 269, 2018 234, 2019 180, 2020 57 | |
| **SI Figure 2b** | <550 449, 550-650 1778, 650-750 545, 750-850 2745, 850-950 860, ≥950 123 | |
| **SI Figure 2c** | <u>SGC</u><br><br>1999 3, 2000 7, 2001 12, 2002 20, 2003 84, 2004 142, 2005 202, 2006 145, 2007 256, 2008 69, 2009 92, 2010 91, 2011 69, 2012 60, 2013 46, 2014 22, 2015 30, 2016 13, 2017 7 | <u>Non-SGC</u><br><br>1988 3, 1989 3, 1990 12, 1991 11, 1992 20, 1993 16, 1994 15, 1995 34, 1996 23, 1997 29, 1998 116, 1999 145, 2000 157, 2001 208, 2002 194, 2003 186, 2004 200, 2005 207, 2006 204, 2007 188, 2008 164, 2009 198, 2010 206, 2011 338, 2012 374, 2013 363, 2014 341, 2015 330, 2016 314, 2017 262, 2018 232, 2019 178, 2020 56 |

**Supplementary Table 4.** List of structural genomic consortia in a dataset of structures not filtered by chemical shift completeness. Note that some structures were jointly produced by multiple SGCs.

| **Structural genomics consortium** | **# of structures** |
|---|---|
| RIKEN Structural Genomics/Proteomics Initiative (RSGI) | 617 |
| Northeast Structural Genomics Consortium (NESG) | 518 |
| Joint Center for Structural Genomics (JCSG) | 84 |
| Center for Eukaryotic Structural Genomics (CESG) | 55 |
| Seattle Structural Genomics Center for Infectious Disease (SSGCID) | 44 |
| Structural Proteomics in Europe (SPINE) | 44 |
| Structural Genomics Consortium (SGC) | 35 |
| Ontario Centre for Structural Proteomics (OCSP) | 34 |
| Membrane Protein Structures by Solution NMR (MPSbyNMR) | 16 |
| New York Structural Genomics Research Consortium (NYSGRC) | 13 |
| Partnership for T-Cell Biology (TCELL) | 13 |
| Berkeley Structural Genomics Center (BSGC) | 10 |
| Structure 2 Function Project (S2F) | 9 |
| Center for Structural Genomics of Infectious Diseases (CSGID) | 7 |
| Montreal-Kingston Bacterial Structural Genomics Initiative (BSGI) | 7 |
| Midwest Center for Structural Genomics (MCSG) | 6 |
| Partnership for Stem Cell Biology (STEMCELL) | 6 |
| Mitochondrial Protein Partnership (MPP) | 6 |
| Chaperone-Enabled Studies of Epigenetic Regulation Enzymes (CEBS) | 5 |
| New York SGX Research Center for Structural Genomics (NYSGXRC) | 4 |
| Center for Structures of Membrane Proteins (CSMP) | 2 |
| TB Structural Genomics Consortium (TBSGC) | 1 |
| Assembly, Dynamics and Evolution of Cell-Cell and Cell-Matrix Adhesions (CELLMAT) | 1 |
| Southeast Collaboratory for Structural Genomics (SECSG) | 1 |

**Supplementary Figure 3.** Dependence of correlation score and RMSD score on other measures of structural accuracy. The number of samples and the mean are indicated below each box. (a) Number of NOE restraints per residue. (b) Number of dihedral restraints per residue. (c) Number of hydrogen bond restraints per residue. (d) Mean size of distance restraint violation (Å). (e) Mean size of dihedral restraint violation (°). (f) Mean backbone root-mean-square-difference (RMSD – the precision). (g) Percentage of backbone (φ, ψ) pairs within the favoured regions of the Ramachandran plot. (h) Clashscore (clashes per 1000 atoms).

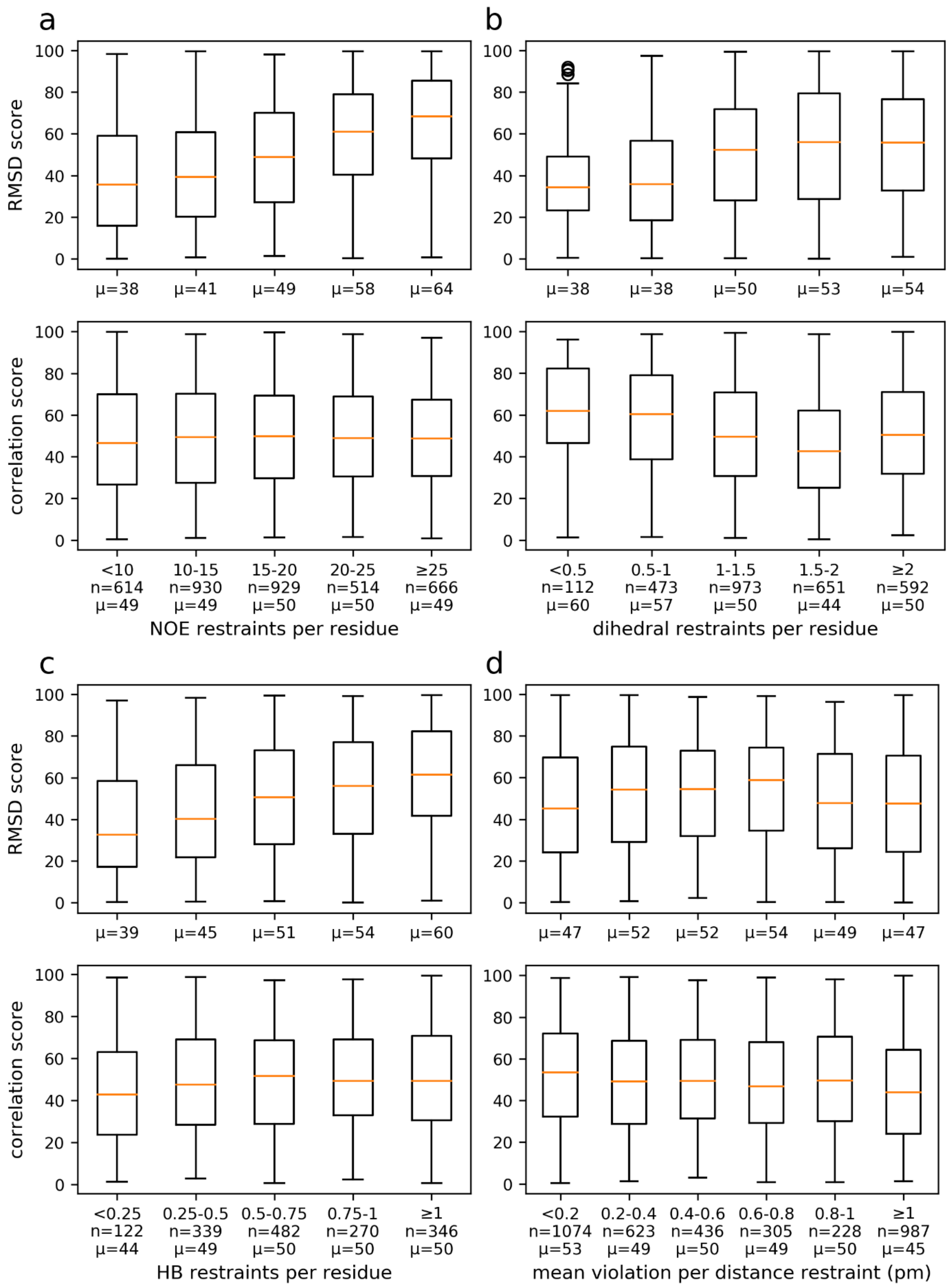

**Supplementary Figure 3 continued.**

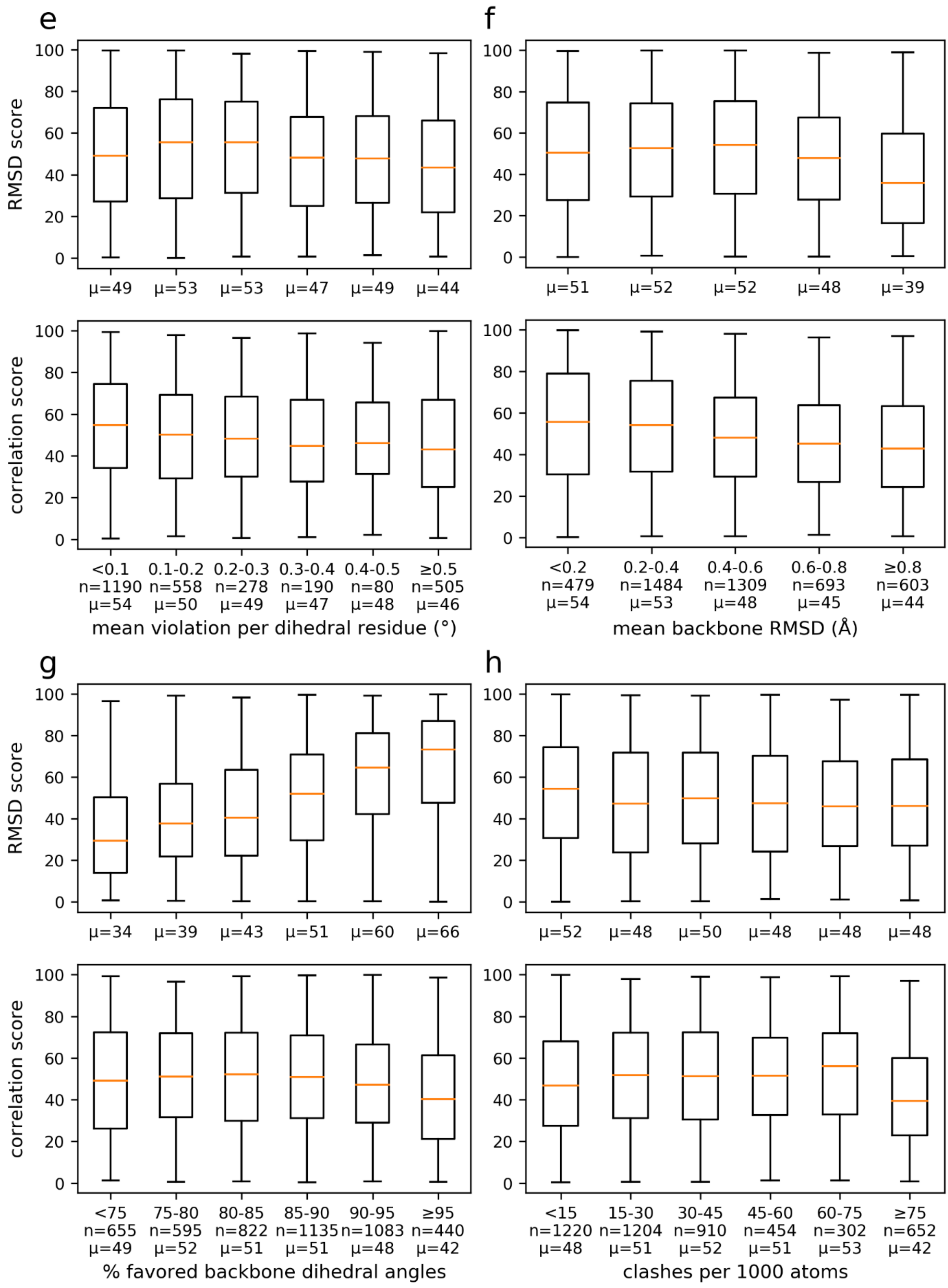
